# Multi-modal analysis of satellite cells reveals early impairments at pre-contractile stages of myogenesis in Duchenne muscular dystrophy

**DOI:** 10.1101/2025.01.24.634739

**Authors:** Sophie Franzmeier, Shounak Chakraborty, Armina Mortazavi, Jan B. Stöckl, Jianfei Jiang, Nicole Pfarr, Benedikt Sabass, Thomas Fröhlich, Clara Kaufhold, Michael Stirm, Eckhard Wolf, Jürgen Schlegel, Kaspar Matiasek

## Abstract

Recent studies on the role of myogenic satellite cells (SC) in Duchenne muscular dystrophy (DMD) documented altered division capacities and impaired regeneration potential of SC in DMD patients and animal models. It remains unknown, however, if SC-intrinsic effects trigger these deficiencies at pre-contractile stages of myogenesis rather than resulting from the pathologic environment. Addressing this, we isolated SC from muscle biopsies of a porcine DMD model for characterization. Traction force microscopy (TFM) revealed that DMD SC produce a significantly higher strain energy than wild-type cells (WT; 0.136 ± 0.016 µJ vs. 0.057 ± 0.008 µJ). By RNA-seq, we identified 1,390 differentially expressed genes and proteomics measurements detected 1,261 proteins with altered abundance in DMD vs. WT. Dysregulated pathways uncovered by Gene Ontology (GO) enrichment analysis included sarcomere organization, focal adhesion, and response to hypoxia. We integrated the data using multi-omics factor analysis (MOFA) and identified five factors accounting for the variance with an overall higher contribution of the transcriptomic (61.95 %) than the proteomic data (54.02 %). Our findings suggest SC impairments result from their inherent genetic abnormality rather than environmental influences. The observed biological changes are independent and not reactive to the pathological surrounding of DMD muscle.

## Introduction

Over the last two decades, studies on Duchenne muscular dystrophy (DMD) focused on the pathology of differentiated, dystrophic muscle fibers while leaving a gap in the comprehension of early disease mechanisms in pre-contractile stages of myogenesis. It was not until the discovery of high expression levels of dystrophin in activated satellite cells (SC) (Dumont et al., 2015), the stem cells of skeletal muscle tissue (Yin et al., 2013), that the interest in the role of SC in the development of DMD grew considerably. On evolving concepts, several studies detected elevated numbers of myogenic SC in muscle fibers of DMD patients (Bankole et al., 2013; Kottlors & Kirschner, 2010) and *mdx* mice (Reimann et al., 2000; Ribeiro et al., 2019) which was explained by the hypothesis of SC exhaustion (Heslop et al., 2000; Luz et al., 2002). Recent investigations related these observations to the impairment of asymmetric divisions leading to an imbalance in maintaining the SC pool through self-renewal and the generation of muscle progenitor cells (Dumont & Rudnicki, 2016; Dumont et al., 2015). Still, it needs to be clarified if these impairments result from the pathological environment SC reside in, or whether there are intrinsic SC-defects, emerging at the undifferentiated muscle progenitor cell state. An investigation of dystrophin-deficient SC and their function on disease onset and progression should lead to a better understanding of DMD pathobiology. As stem cell-based approaches may offer promising therapy options (Sun et al., 2020), the elucidation of affected biological pathways in DMD SC is an urgent research topic.

First described in 1961 by Alexander (Mauro), SC are indispensable for muscle growth, regeneration, and repair. Located between the basal lamina and the sarcolemma, they reside in a quiescent state and are only activated upon muscle trauma or injury. Once stimulated, SC proliferate extensively and by undergoing asymmetric division, they give rise to myogenic precursor cells, which enter the myogenic pathway to differentiate into myofibers while simultaneously maintaining the stem cell pool through self-renewal (Relaix et al., 2021; Yin et al., 2013). The steps of myogenesis are determined by the expression of the myogenic transcription factors PAX7 (paired box protein 7), (Buckingham & Relaix, 2015), MYF5 (myogenic regulatory factor 5), MYOD1 (myoblast determination protein 1), and MYOG (muscle-specific transcription factor myogenin) (Zammit, 2017). Any disturbances to this precisely paced process have substantial effects on muscle homeostasis; consequently, SC dysfunction is supposed to play a crucial role in the progression of muscle pathology and failed regeneration in various myopathies (Ganassi et al., 2022; Ganassi & Zammit, 2022).

DMD is one of the most devastating muscle dystrophies in people, an X-linked disease affecting approximately one in 3500 – 5000 male births and ultimately leading to death in early adulthood (Emery, 1991; Happi Mbakam & Tremblay, 2023; Mendell et al., 2012). It is caused by mutations in *DMD*, the largest known gene in the human genome spanning 2.5 Mb and harboring 79 exons. *DMD* encodes dystrophin, an actin-binding structure protein primarily expressed in skeletal and heart tissue where it constitutes part of the dystrophin-associated glycoprotein complex (DGC) (Gao & McNally, 2015). The DGC connects cytoskeletal F-actin filaments of the myofiber through the sarcolemma to the extracellular matrix (ECM) and is responsible for muscle fiber integrity and stability. In humans, *DMD* mutations occur mainly in hotspot regions between exons 2 - 10 and 45 - 55. These mutations are most frequently deletions of one or several complete exons (∼°60 - 70 %), followed by point mutations (∼°20%) and exon duplications (∼°5 - 15%) leading to a reading frame shift which ablates dystrophin expression (Duan et al., 2021). Lack of dystrophin causes instability of the sarcolemma with a rapidly progressing fiber degeneration, which severely compromises the contractile apparatus of the affected muscle tissue (Fontelonga et al., 2019). Starting at a young age, affected males show a rapidly progressive muscle weakness leading to walking disabilities and patients decease in their 2^nd^ or 3^rd^ decade of life due to respiratory and cardiac failure (Passamano et al., 2012; Wahlgren et al., 2022).

Currently, there are almost 60 different animal models available for DMD research, including mammals (e.g. *Mus musculus*) and non-mammals (e.g. *Caenorhabditis elegans*, *Danio rerio*) (McGreevy et al., 2015; Zaynitdinova et al., 2021) but when it comes to translational studies, large animal models, reflecting the human conditions most accurately, are of urgent need. On that note, aground great similarities on an anatomical and physiological level to humans, the pig as a pre-clinical model for DMD research has attracted consideration for medical research (Moretti et al., 2020; Stirm et al., 2021; Stirm et al., 2024). The first porcine DMD model was developed in 2013 and harbors a DMD exon 52 deletion, which leads to a disease phenotype similar to human patients (Klymiuk et al., 2013). These models are predominately used for investigations of differentiated, contractile myofibers but offer a great opportunity to expand their application toward research exploring the behavior of dystrophic muscle stem cells.

Motivated by these challenges we used fresh muscle samples from dystrophin-deficient DMD and wild-type (WT) control piglets and optimized current protocols of SC isolation for use in porcine tissue. Then, free from the influence of the dystrophic environment, enabling the observation of intrinsic cellular changes, we conducted a comprehensive characterization of the isolated SC. To monitor different developmental stages of myogenesis, we analyzed cultivated SC during proliferation and after induction to differentiate into multinucleated myotubes. After assessing growth capacities and force generation *in vitro*, we performed bulk RNA sequencing and label-free proteomics to screen for molecular derangements associated with dystrophin-deficiency. To find sources explaining the observed variance, we utilized multi-omics factor analysis (MOFA) to integrate the transcriptomic and proteomic datasets. We detected increased force generation as well as significantly altered transcriptome and proteome profiles at early and late stages of myogenesis, which unambiguously could be assigned to the lack of dystrophin. Our data strongly suggest that dystrophin deficiency indeed highly affects the intrinsic behavior of muscle SC and impacts myocyte differentiation already at a pre-contractile stage.

## Results

### 1. Muscle sections of DMD animals harbor significantly elevated levels of PAX7+ SC

To verify dystrophin-deficiency in DMD tissue, we conducted an immunohistochemical (IHC) staining assay against dystrophin on muscle sections derived from *intercostalis (INT)* and *pectoralis (PECT)* muscle samples of male DMD (n=6) and WT (n=6) animals. As anticipated, dystrophin signal was absent in DMD tissue, whereas in WT a strong sarcolemma-associated signal was observed. There was no obvious difference in dystrophin expression between *INT* and *PECT* sections (**Fig. 1A**).

**Figure 1:**
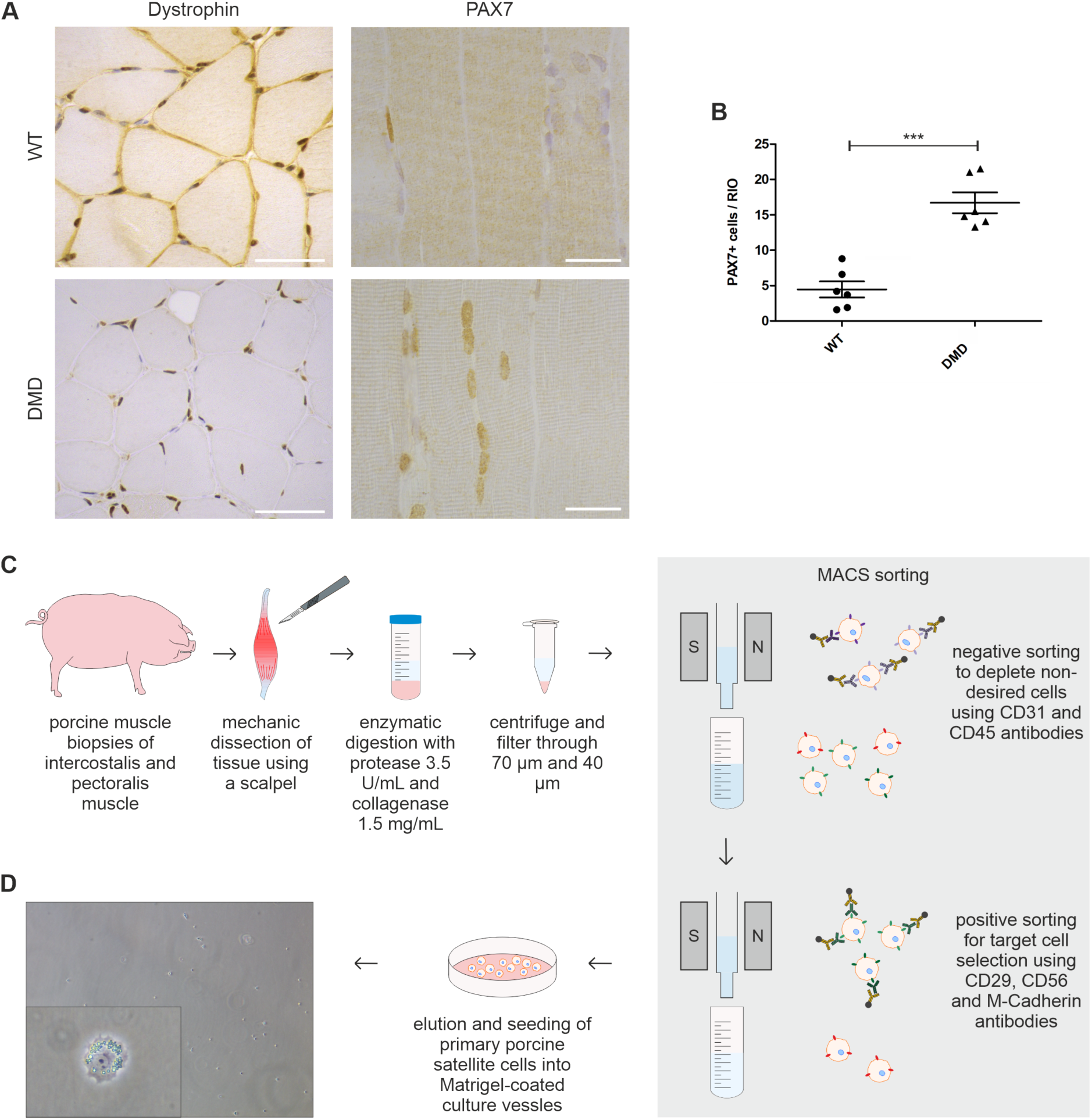
Detection of PAX7+ SC in dystrophin-deficient muscle sections and SC isolation pipeline. **(A)** Immunohistochemistry of dystrophin and PAX7 in paraffin-embedded PECT muscle sections of DMD and age-matched WT controls. DMD tissue demonstrates absent sarcolemmal dystrophin signal (brown) and high amounts of PAX7+ cell nuclei (brown). Scale bars = 50 µm (dystrophin) and 25 µm (PAX7). **(B)** Significantly higher amounts of PAX7+ cells were detected in DMD (n=6, triangle, 16.70 ± 1.471) muscle sections when compared to WT (n=6, dots, 4.467 ± 1.137; p=0.0001); each dot represents one individual; unpaired t-test was used for statistical testing and results are shown as mean and SEM. **(C)** Schematic overview of isolation procedure of primary porcine SC derived from INT and PECT muscle biopsies of DMD and WT animals. **(D)** Representative light microscopy pictures of seeded SC directly after isolation; insert showing magnetically labeled cell. Magnification 10x and 40x respectively.

To identify SC and compare the size of the residing stem cell population in muscle sections of DMD (n=6) and WT piglets (n=6), we performed an immunohistochemistry (IHC) staining against PAX7, a widely used marker for quiescent and early activated SC (Ropka-Molik et al., 2011; Zammit et al., 2006) (**Fig. 1A**). We detected PAX7+ cells at the expected location at the stem cell niche between the sarcolemma and the basal lamina of the muscle fascicle. Quantification of IHC sections revealed that the amount of PAX7+ cells in *INT* muscle of DMD (16.70 ± 1.471) was 3.7-times higher compared to WT pigs (4.467 ± 1.137; p=0.0001) (**Fig. 1B**). Regarding *PECT*, similar results were obtained with a likewise significantly increased number of PAX7+ cells in DMD pigs compared to WT controls (data not shown).

These results are in line with previous studies detecting higher numbers of PAX7+ SC in muscle tissue of young *mdx* mice, a murine model for DMD (Jiang et al., 2014; Kottlors & Kirschner, 2010; Ribeiro et al., 2019). Further, we demonstrate that the increase in PAX7+ SC is similar in both muscles involved in respiration and movement.

### 2. Isolated PAX7+ SC from DMD piglets lack dystrophin expression

To accurately isolate SC from porcine skeletal muscle biopsies, we developed a modified isolation protocol using a two-step magnetically activated cell sorting (MACS) system with antibodies to deplete undesired cells and to specifically target SC simultaneously (**Fig. 1C**). A detailed version of our protocol is provided in **Supplemental Methods**. Efficacy of the protocol in terms of cell yield and purity of the population was assured by immunofluorescence (IF) staining of cultured cells against desmin, a marker for skeletal muscle intermediate filaments (Paulin & Li, 2004; Yang et al., 2017) and against PAX7. In all samples across the two genotypes, isolated cells presented expression of both makers, confirming the selective isolation of skeletal muscle stem cells (**Fig. 2A**). To evaluate dystrophin-deficiency in the isolated cells, we conducted a Western blot analysis using antibodies against the rod domain (DYS1) and the C-terminus (DYS2) of dystrophin. WT SC showed a clear band at approximately 427 kDa corresponding to the full-length muscle isoform of dystrophin as well as one smaller isoform at 268 kDa. In contrast, dystrophin expression was absent in DMD SC (**Fig. 2B**).

**Figure 2:**
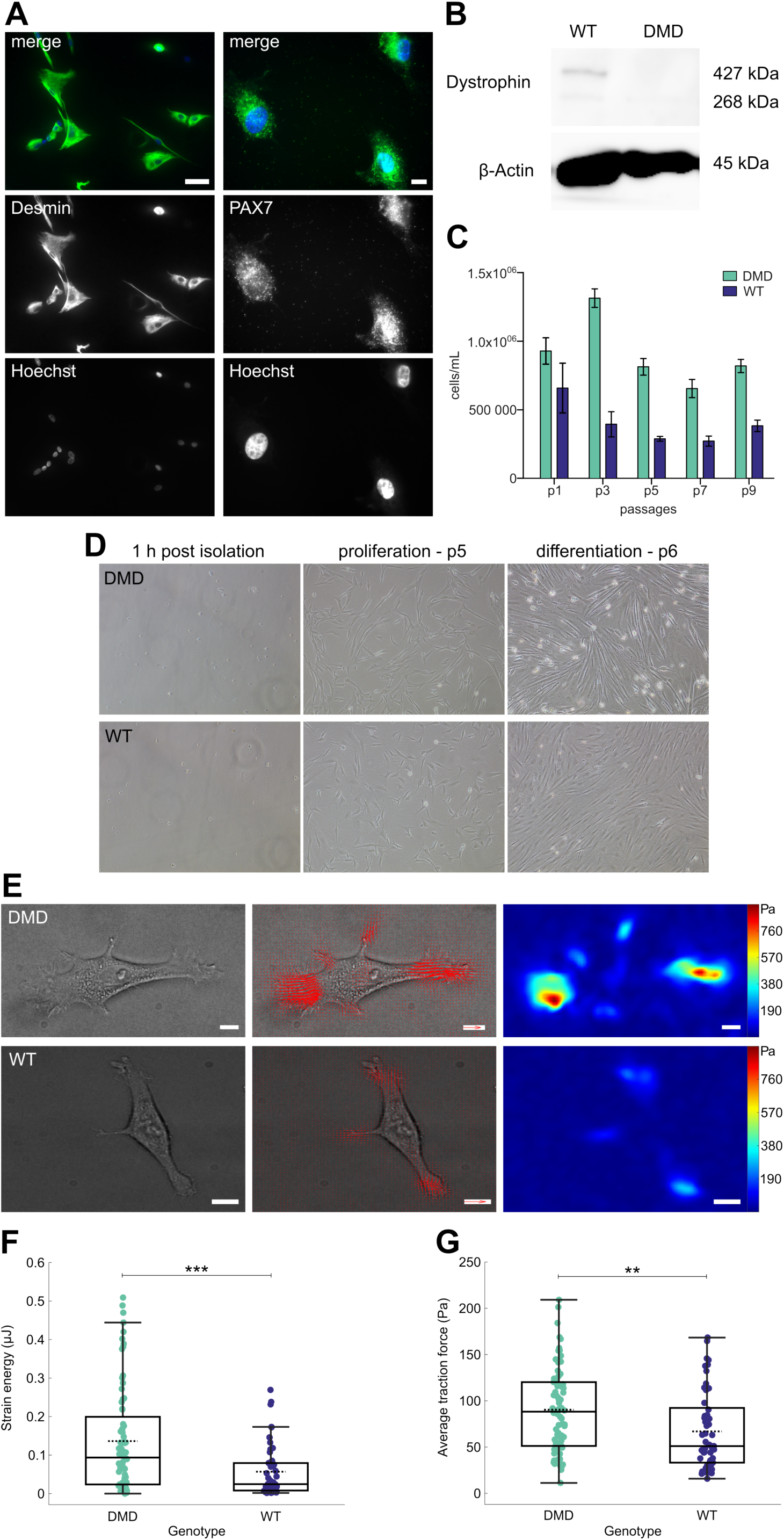
Characterisation of isolated primary porcine SC and quantification of traction forces in single DMD and WT SC. **(A)** Immunofluorescence staining of cultured WT SC after isolation using desmin (green) and PAX7 (green) antibodies to confirm the cells’ identity. Hoechst (blue) was used as a nuclear counterstain. Magnification desmin 20x, scale bar = 50 µm. Magnification PAX7 63x, scale bar = 30 µm. **(B)** Western blot analysis to detect dystrophin expression in DMD and WT SC lysates. A band at approx. 427 kDa indicates dystrophin expression in WT cells which is completely absent in lysates of DMD cells. β-Actin was used as loading control. **(C)** Amount of cells/mL for DMD (n=4, green) and WT (n=4, purple) over the span of 9 passages (p) in vitro, values are displayed as mean and SEM. **(D)** Representative bright field pictures of primary porcine SC culture in vitro 1h post isolation, during proliferation at p5 and after serum withdrawal to induce differentiation at p6. Magnification 10x. **(E)** Representative bright field images of DMD and WT SC (left) with corresponding quiver plots (middle) and traction maps (right); scale bar = 10 µm; traction vector (red quiver) 1 kPa. **(F)** Strain energy (DMD, green, 72 cells, 0.136 ± 0.016 µJ; WT, purple, 50 cells, 0.057 ± 0.008 µJ, p=0.00052) and **(G)** average traction force levels (DMD 90.4 ± 10.5 Pa; WT 66.9 ± 8.9 Pa; p=0.0018) visualized as box plots overlaid with swam charts; each dots represents one cell; two-tailored Mann-Whitney U test was used for statistical testing and results are presented as mean ± SEM.

These data show that our protocol allows the isolation of porcine SC and lend proof of failed dystrophin expression in DMD SC, thus we concluded to use the cells for all subsequent experiments in this study.

### 3. DMD SC morphology under proliferation and differentiation conditions

Following MACS isolation, SC were seeded into Matrigel-coated culture vessels and SC from both DMD and WT animals uniformly started to adhere within 12 h (**Fig. 1D**). In this period the cells of both genotypes displayed a small, round shape with approximately 5 – 10 µm diameter. Upon adherence, the cells successively increased in length and volume and displayed a clear cytoplasm with a prominent nucleus. After 3 – 4 days in culture, the cells presented with typical arborized, stellate myoblast morphology with several extending processes (**Fig. 2D**). Proliferating SC were cultivated in standard growth medium until reaching °70 - 80% confluence. Thereafter, the cells were passaged every other day and manually counted to estimate cell numbers. Throughout 9 passages (p), we observed differences between cell numbers of DMD (n=4) and WT SC (n=4). DMD numbers peaked at p3 and declined slightly until p9. In contrast, cell numbers of WT SC were highest at p1, then steadily decreased and maintained roughly the same (**Fig. 2C**). To induce differentiation in DMD and WT SC, we performed serum deprivation for 10 days. On day 4 of serum reduction, the cells of both groups increased in length and displayed an elongated, bipolar shape. Aligned, multinucleated myotubes were observed on day 6 (**Fig. 2D**). For the following days, the myotubes continued to grow in size and further fused. We did not observe any major differences between DMD and WT myotubes.

### 4. DMD SC exhibit increased force generation

To investigate how dystrophin-deficiency affects force generation and contractility in SC, we employed traction force microscopy (TFM). We seeded the cells onto polydimethylsiloxane (PDMS) substrates with a Young’s modulus of approx. 12 kPa, which closely mimics the physiological stiffness of muscle tissue (Bruyere et al., 2019). A single layer of fluorescent nanospheres was embedded on the substrate surface to enable precise force measurements. To quantify the magnitude of the mechanical forces that cells exert on their surrounding environment, we measured the displacement of the beads within the substrate and subsequently calculated traction forces fields using standard Bayesian Fourier Transform Traction Cytometry (**Fig. 2E**). We observed that the strain energies generated by individual cells were, on average, significantly higher for DMD SC (n=4, 72 cells overall, 0.136 ± 0.016 µJ) than for WT SC (n=4, 50 cells overall, 0.057 ± 0.008 µJ; p=0.00052) (**Fig. 2F**). Additionally, exerted average traction forces were significantly higher for DMD (90.4 ± 10.5 Pa) than for WT SC (66.9 ± 8.9 Pa; p=0.0018) (**Fig. 2G**).

Thus, dystrophin-deficiency in muscle SC influences force generation already at a level of individual cells, which highlights its significance to the pathological characteristics of DMD, which in turn could affect cell-ECM interactions and mechanotransduction (Gao & McNally, 2015).

### 5. Overview of transcriptome profiles of DMD and WT SC

To obtain insight into the effects of dystrophin-deficiency on the transcriptome of SC in a proliferative state (PROL) as well as during differentiation into myotubes (DIFF), RNA isolated from DMD (n=4) and WT (n=4) SC samples was analyzed by 3’ mRNA-seq (adapted from Bagnoli et al., 2018; Picelli et al., 2013; Picelli et al., 2014). Principal component analysis (PCA) clearly separated the transcriptome profiles of DMD SC and WT SC, both at the PROL and DIFF states, in the first two principal components (PC1 and PC2) (**Fig. 3A**). Unsupervised hierarchical clustering further confirmed the genotype- and state-dependent separation of transcriptome profiles (**Fig. 3B**).

**Figure 3:**
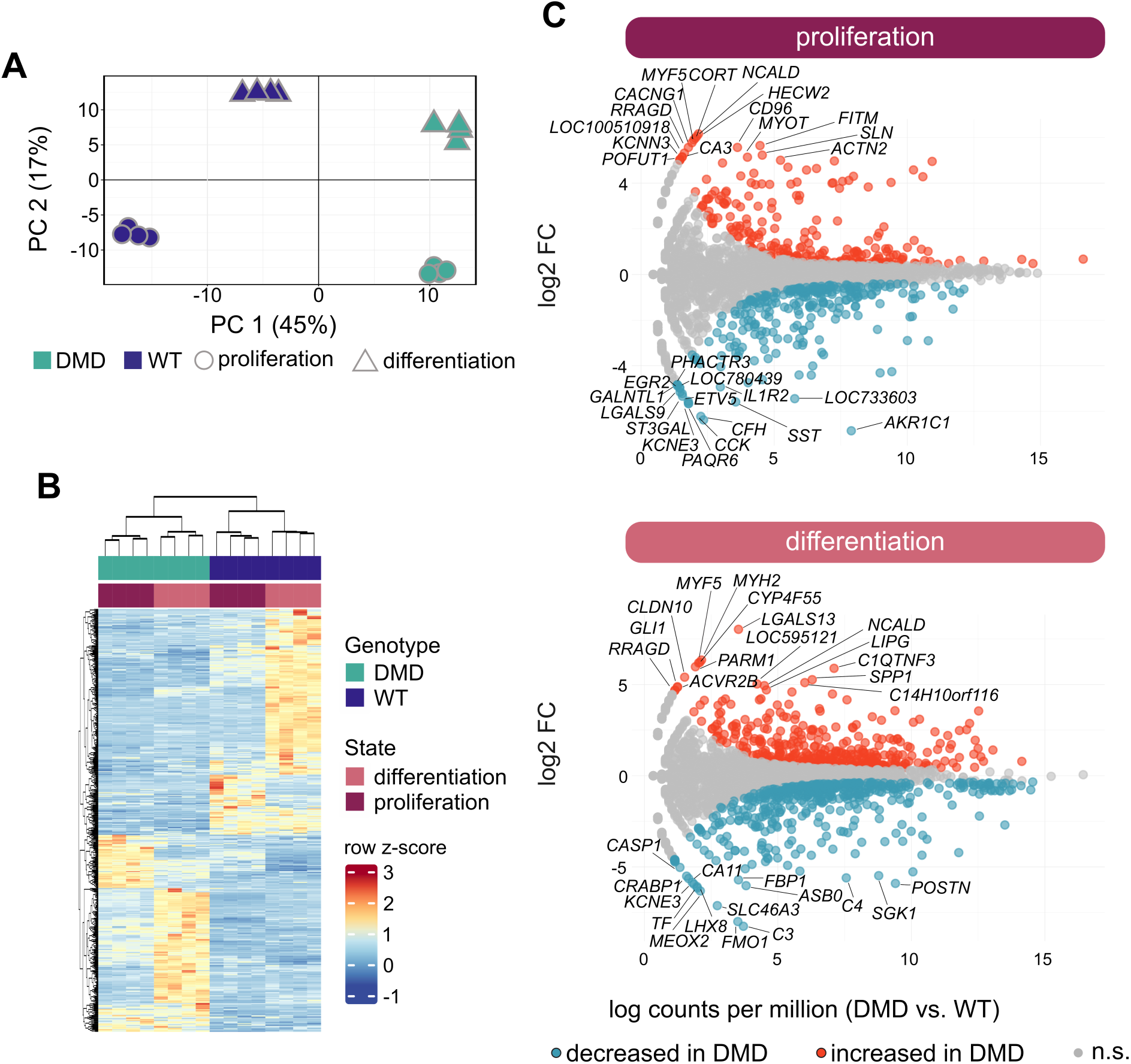
Transcriptome analysis of DMD and WT SC. **(A)** Principal component analysis of the transcriptome profiles of SC derived from DMD (n=4, green) and WT (n=4, purple) piglets in proliferation (dots) and differentiation (triangle) state; each symbol represents one transcriptome profile. The percentage of variation of each component is indicated. **(B)** Heat map generated by unsupervised hierarchical clustering of top 500 differentially expressed genes between DMD and WT SC. **(C)** Total number of differentially expressed genes between DMD and WT (adjusted p-value < 0.05, |log2 fold change| > 0.6) displayed as MA plot for proliferation **(D)** and differentiation **(E)** state, red color indicates up- and blue color downregulated genes in DMD SC respectively, genes are represented as dots.

These findings emphasize that dystrophin-deficiency could indirectly play a role in the regulation of gene expression in yet undifferentiated states of muscle development.

### 6. Differences in the transcriptome of DMD vs. WT SC in a proliferative respectively differentiated state

In the PROL state, we identified 568 differentially expressed genes (adjusted p-value < 0.05; |log2 fold change| > 0.6) between DMD and WT SC (261 with higher and 307 with lower expression levels in DMD). Amongst genes showing the strongest upregulation in DMD, we detected *HECW2*, *NCALD*, *CORT,* and several key myogenesis genes including *MYF5*, *MYOD1,* and *MYOG*. The top-downregulated genes included *AKR1C1*, *CFH, CCK,* and *PAQR6* (**Fig. 3C**). Using the DAVID online tool, we conducted a gene ontology (GO) enrichment analysis using the biological process (BP) database. Upregulated genes in DMD SC showed a significant enrichment of gene sets related to skeletal muscle fiber development, sarcomere organization, skeletal muscle contraction, and positive regulation of myoblast differentiation. On the other hand, the top-downregulated genes in DMD SC were associated with the ontologies of inflammatory response, positive regulation of angiogenesis, response to hypoxia and cell migration, and adhesion.

Analyzing the DIFF state, we detected 822 differentially expressed genes (adjusted p-value < 0.05; |log2-fold change| > 0.6) between DMD and WT SC (391 with higher and 431 with lower expression levels in DMD). Genes displaying the strongest increase in expression in DMD SC were *LGALS13*, *MYF5*, and *MYH2.* Additionally, the mRNA levels of smooth muscle related genes like *TAGLN* and *CNN1* as well as myosin light chain genes *MYLPF* and *MYL1* were increased. The latter two were also strongly increased in the PROL state (see above). Interestingly, elevated transcript levels of the myogenic transcription factors *MYF5* and *MYOG* were identified. The top-downregulated genes in DMD SC comprise *C3*, *FMO1*, and *SLC46A3*. (**Fig. 3C)**. The GO-BP analysis revealed that the top-upregulated genes in DMD SC displayed a significant enrichment of gene sets associated with muscle contraction, apoptotic process, and glycolytic metabolism. Amongst the top-downregulated genes in DMD SC, we identified significant enrichment of gene sets related to the ontologies of cytoplasmic translation, negative regulation of cell growth, response to oxidative stress, and TGF-β signaling pathway. A detailed list with differentially expressed genes and identified pathways is provided in **Supplemental Table S1 - S4**.

Collectively, these data highlight the diverse role of dystrophin at different stages of myogenesis and show that dystrophin-deficiency affects a broad spectrum of intracellular processes ranging from myoblast-specific pathways to universal cell signaling cascades.

### 7. Overview of the proteome profiles of DMD and WT SC

To achieve a comprehensive overview of the consequences of dystrophin-deficiency in SC, we performed quantitative proteomics using liquid chromatography-mass spectrometry (LC-MS/MS) with DMD (n=4) and WT SC (n=4) in PROL and DIFF states aiming to explore any molecular alterations on the protein level between the genotypes in the different developmental stages.

In total 49115 peptides were identified which could be assigned to 4297 proteins at a false discovery rate (FDR) of 1%. In all samples, characteristic markers for skeletal muscle tissue were detected, namely sarcomere-associated myosin heavy and light chain proteins (e.g. MYH9, MYL6, MYH3) and actins (e.g. ACTG1, ACTA2), contraction-related tropomyosins (e.g. TPM1, TPM4, TPM3) as well as the muscle-specific intermediate filament desmin (DES). A list with all identified and quantified proteins can be found in the **Supplemental Table S5** and **S6**.

The dimensionality of the proteomic data was reduced using PCA, showing a separation of the proteome profiles of DMD and WT SC in both the PROL and DIFF cell states (**Fig. 4A**), which is in line with the findings of the transcriptome analysis. Notably, unsupervised hierarchical clustering of DMD and WT SC in DIFF and PROL states did not separate the groups according to genotype and state but revealed two major clusters. One cluster consisting of only WT DIFF and a second cluster composed of WT PROL, DMD PROL, and DMD DIFF (**Fig. 4B**). This finding demonstrates that DMD SC proteomes are more similar to WT PROL than WT DIFF, indicating impaired differentiation in DMD SC.

**Figure 4:**
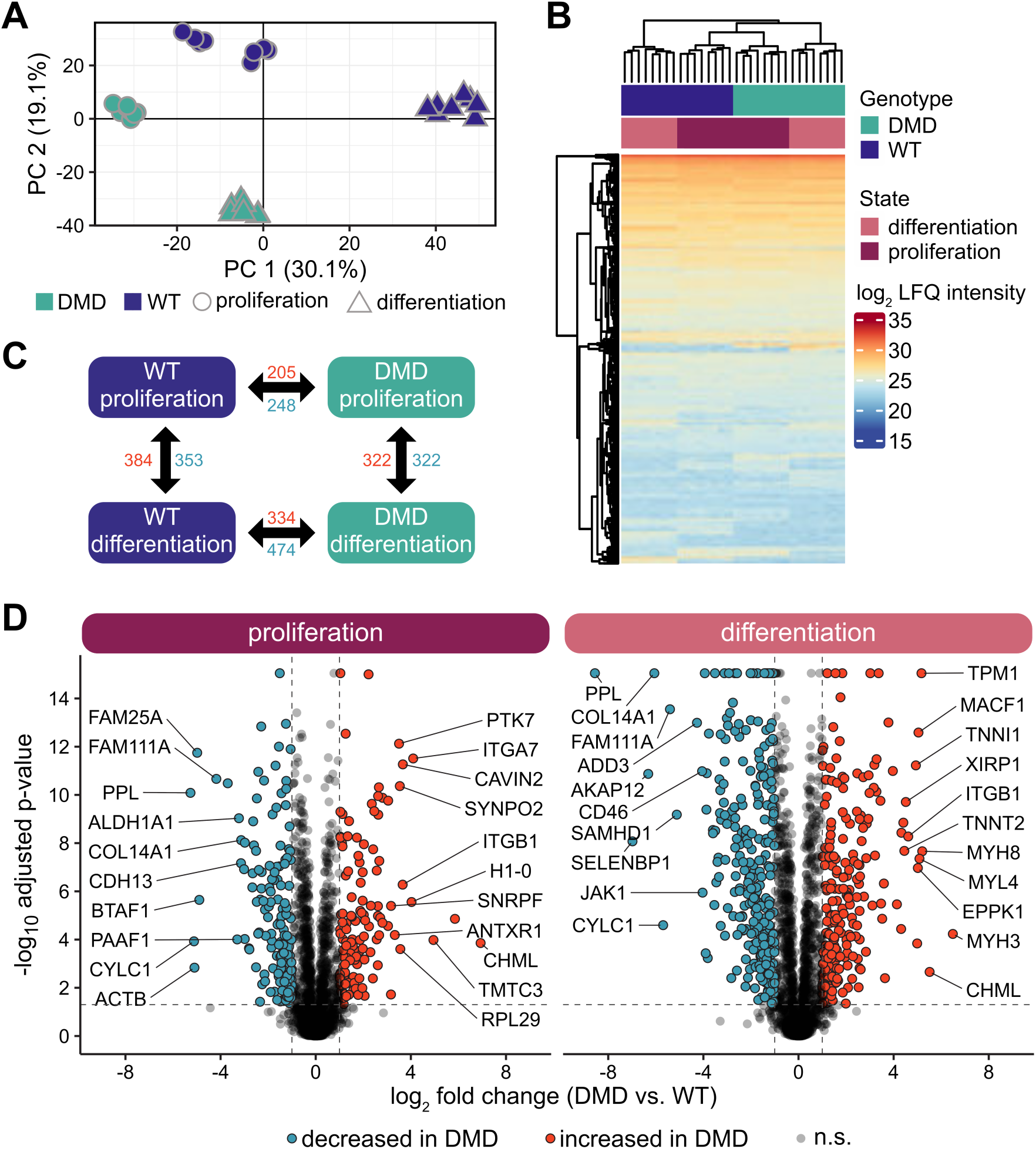
Proteome analysis of DMD and WT SC. **(A)** Principal component analysis of the proteome profiles of SC derived from DMD (n=4, green) and WT (n=4, purple) animals in proliferation (dots) and differentiation (triangle) state; each dot represents one proteome profile. The percentage of variation of each component is indicated. **(B)** Heat map and unsupervised hierarchical clustering of protein intensity values from proteome expression profiles of DMD and WT SC. **(C)** Differentially abundant proteins within (horizontal) and between (vertical) DMD and WT SC proliferation (vertical) and differentiation (horizontal) state. Numbers of proteins with significantly higher (red) and lower (blue) abundance with an adjusted p-value of < 0.05 and a |log2 fold change| > 0.6. **(D)** Volcano plots of differentially abundant proteins between DMD and WT SC in proliferative respectively differentiated state - Two-way ANOVA followed by Tukey Honest Significant Differences was used for statistical testing, as significance cut off adjusted p-value < 0.05 and |log2 fold change| > 0.6 was used. Proteins are represented as dots; red color indicates higher and blue lower abundance in DMD cells.

Taken together, proteomic profiling of DMD and WT SC highlights the complexity of DMD myopathology and points towards a highly altered molecular profile of DMD SC along the myogenic pathway.

### 8. Differences in the proteomes of DMD and WT SC in a proliferative respectively differentiated state

By further statistical analysis, we detected proteins with significantly altered abundances between DMD and WT SC (adjusted p-value < 0.05; |log2 fold change| > 0.6). Analyzing DMD and WT SC in PROL state we identified 453 proteins significantly altered in abundance (205 with increased and 248 with decreased abundance in the DMD samples). Amongst proteins with the highest increase in abundance, we found ITGA7, ITGB1, DES, and TPM1 whereas among the strongest decrease in abundance proteins included PPL, ACTB, and MYH11. In DIFF state the number was even higher with 808 differentially abundant proteins (334 with increased and 474 with decreased abundance in the DMD samples) (**Fig. 4C**). Top differentially expressed proteins included several myosins (MYH3, MYH8, MYL4, MYL11), troponins (TNNI1, TNNT2, TNNC1), and MACF1 with higher and PPL, CYLC1, ALDH1A3, and ALDH1A1 with lower abundance.

Using DAVID on the differentially abundant proteins, we investigated several functional GO categories such as biological process, cellular component, molecular function as well as the Kyoto Encyclopedia of Genes and Genomes (KEGG) database. In PROL, clusters of proteins with increased abundance in DMD compared to WT showed significant enrichment in proteins related to mRNA splicing ribosome/translation, structural constituent of cytoskeleton, DNA replication, and focal adhesion. Sets of proteins with decreased abundance in DMD were related to GTP binding, biosynthesis of nucleotide sugars, and regulation of cytoskeleton. At the DIFF state, proteins with higher abundance in DMD SC were associated with the regulation of actin cytoskeleton, ribosome, hypertrophic cardiomyopathy, and pathways of neurodegeneration. Analyzing proteins of lower abundance in DMD SC, we detected a significant enrichment in protein sets corresponding to GTPase activity, biosynthesis of nucleotide sugars, microtubule cytoskeleton, proteoglycans in cancer, and aldehyde dehydrogenase activity. Changes in protein abundance are displayed as Volcano plots in **Fig. 4D**. A detailed list of differentially abundant proteins can be found in **Supplemental Table S7** and **S8**.

Apart from changes caused by the lack of dystrophin between the genotypes, we found state-dependent alterations within the genotype. In DMD SC, 578 proteins were significantly altered in abundance (256 with increased and 322 with decreased abundance) in DIFF compared to PROL state, and in WT SC 737 proteins were differentially abundant (384 with increased and 353 with decreased abundance) in DIFF compared to PROL state (**Fig. 4C**).

In conclusion, we discovered various new factors that are subject to regulation by dystrophin-deficiency on a protein level, and our data further reiterated the high degree of proteome alterations in dystrophin-mutated individuals.

### 9. Integration of transcriptomic and proteomic data

In our study, we used MOFA on the transcriptome and proteome data set to find factors that account for the variability between DMD and WT SC and characterized the features that contribute to these factors. The cumulative contribution to the variance was 61.95 % from the transcriptomic and 54.02 % from the proteomic data set (**Fig. 5A**). We identified five factors explaining the variance between DMD and WT SC. Within these factors, factors 1 and 2 explain the highest percentage of variance in both –omics datasets, with factor 1 accounting for 31.33 % and 32.30 % and factor 2 for 23.63 % and 15.03 % in gene and protein expression variability respectively (**Fig. 5B**). Factor 1 discriminated the WT PROL and DIFF from DMD PROL and DIFF while factor 2 separated the samples into WT DIFF and DMD DIFF apart from WT PROL and DMD PROL (**Fig. 5C**). The top 25 features for factor 1 and 2 in transcriptomic and proteomic modality with their sample expression can be found in **Supplemental Figure S1**. Heat maps for factor 1, which distinguished DMD from WT SC, showed in both –omics modalities two major clusters: one consisting of only WT DIFF and a second one with DMD PROL, DMD DIFF, and WT PROL (**Fig. 5D-G**). This finding recapitulates our previous observation on the proteomic profiles of the DMD and WT SC. Heat maps for factor 2, which separated PROL from DIFF cells, revealed that this separation is more driven by the transcriptomic dataset whereas in the proteomic view the DMD DIFF samples clustered apart from the rest of the samples (**Fig 5C-D**). When performing GO enrichment analysis for the top positive factor 1 weights (more abundant in WT), enriched pathways were autophagy, immune response, response to hypoxia, and cell differentiation in the transcriptome data while protein transport, ER to Golgi vesicle-mediated transport and actin cytoskeleton organization were enriched in the proteome data. Top negative factor 1 weights (more abundant DMD) were associated with cell cycle and division, sarcomere organization, muscle contraction, and homeostasis in the transcriptome data and translation, mRNA splicing, and ribosome assembly in the proteome data. GO enrichment analysis for top positive factor 2 weights (more abundant in DIFF) in the transcriptome revealed processes related to metabolism, G protein signaling, and response to insulin while in the proteome layer pathways of fatty acid biosynthesis, mitochondrial ATP synthesis and response to oxidative stress were enriched. Top negative factor 2 weights (more abundant in PROL) were associated with protein folding, translation, and cell cycle transition in both -omics modalities.

**Figure 5:**
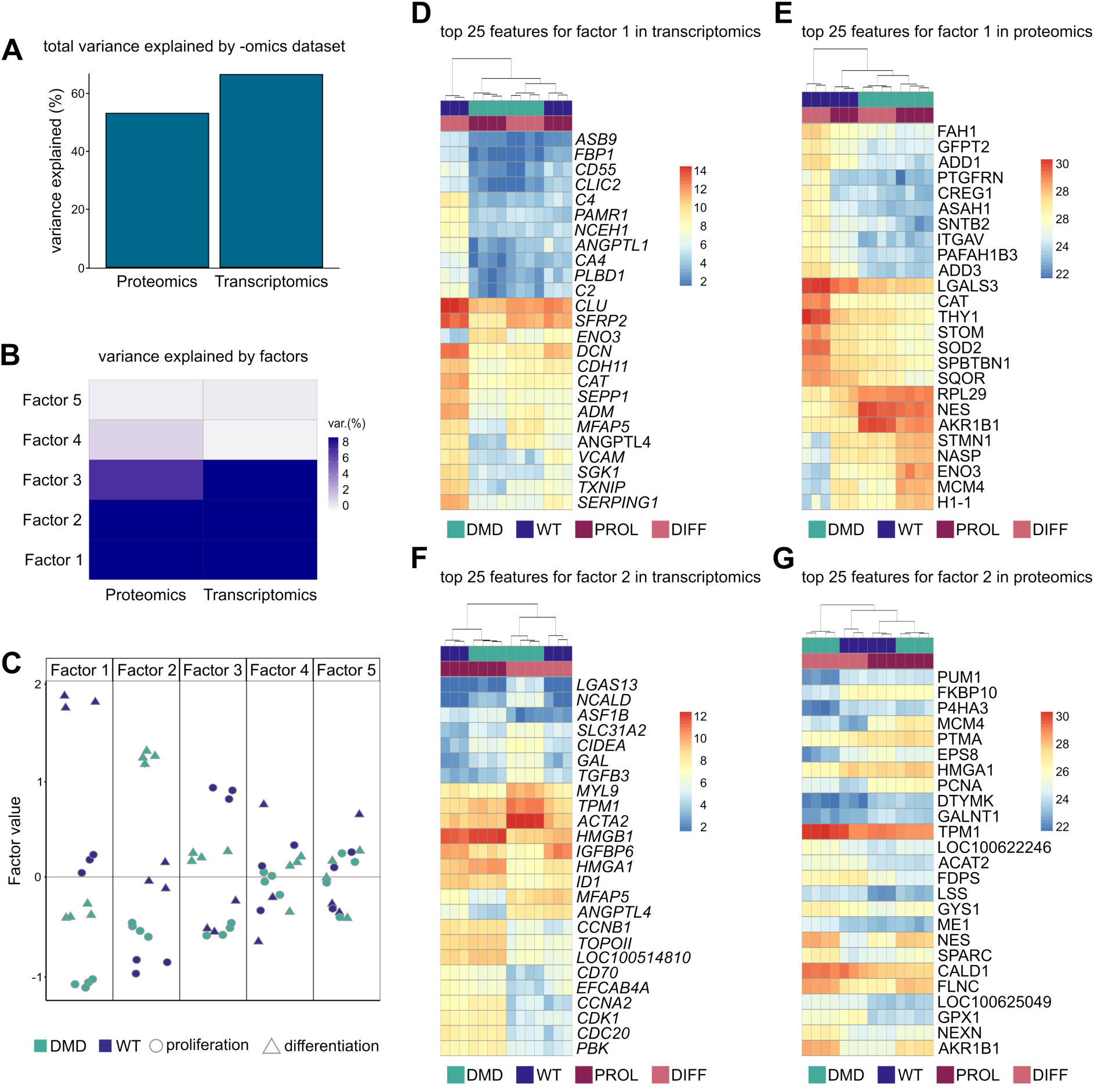
Multi-omics factor analysis of transcriptome and proteome of DMD and WT SC. **(A)** Cumulative variance explained by each -omics modality in percentage. **(B)** Percentage of variance explained by factors identified by multi-omics factor analysis within the transcriptomic respectively proteomic data. Higher percentages of variance among the –omics layers indicate a higher amount of shared variability. **(C)** Swam plot of factor values reveals clustering of DMD (n=4, green) and WT SC (n=3, purple) in proliferation (dots) and differentiation (triangle); each dot represents one individual. **(D)** Top 25 features contributing to Factor 1 from transcriptomic and **(E)** proteomic data. **(F)** Top 25 features contributing to Factor 2 from transcriptomic and **(G)** proteomic data. Heat maps display the distribution of the features across the samples.

## Discussion

For the last two decades, tissue research in DMD focused on the pathology of the mature myofiber, investigating sarcolemma weakening, myofiber necrosis, muscle atrophy, myofibrosis, and inflammation in contractile stages of muscle development (Chang et al., 2016). The discovery of dystrophin expression in murine SC (Dumont et al., 2015) provided new insights into the pathophysiological of DMD and recent studies explored the association of diminished regenerative capacity in dystrophin-deficient SC. However, the origin of these deficiencies remains unclear, in particular, it is still not fully understood if these deficits are a result of the surrounding pathologic stem cell niche or if they emerge from intrinsic cell properties.

In this study, we showed that isolated dystrophin-deficient SC of a porcine model of DMD display profound differences in their gene and protein expression profile in pre-contractile stages of myogenesis. We detected dysregulation of major myogenic transcription factors, indicating a disrupted regeneration capacity in dystrophin-deficient SC that subsequently could contribute to the impaired muscle repair observed in DMD pathology. Additionally, we monitored higher traction forces exerted by the DMD SC, hinting towards modified stiffness and mechanotransduction causing altered cellular responses to external stimuli, which in turn could impact internal signaling pathways.

Due to great similarities with human anatomy, physiology, and metabolism (Zettler et al., 2020) the use of porcine models to generate clinically relevant data is particularly favorable. The DMD pig represents a suitable model for DMD research reflecting the main pathologic features observed in human patients and has greatly enriched disease understanding (Stirm et al., 2022). Thus, we concluded using porcine DMD muscle biopsies for SC isolation offers a great possibility to investigate intracellular effects of dystrophin-deficiency in SC pathogenesis unaffected by the dystrophic niche. Despite higher costs, more intensive care, and stricter ethical regulations compared to rodent models (Burmeister et al., 2022; Hou et al., 2022), porcine tissue offers the advantage of generating large pools of SC, thereby reducing the number of animals needed. In contrast to human samples, porcine muscle biopsies are accessible at young ages, including fetal stages, which allows monitoring of early – even pre-netal – disease events. Standardized procedures and systematic sampling from homogenous age- and/or stage-matched groups of DMD-affected animals ensure sample quality, which is difficult to achieve in human patients’ samples. Cultivating SC *in vitro* additionally offers the chance to monitor DMD implications along the myogenic pathway since SC can be induced to differentiation in a controlled and reproducible manner (Dan-Jumbo et al., 2024; Syverud et al., 2014). On the other hand, phenotypical changes over the time of cultivation are a non-negligible disadvantage of primary cell culture (Vacanti et al., 2005). Therefore, we used only cells until passage number p7 for our experiments. The use of SC offers a feasible system for basic research experiments and genetic manipulations for preclinical studies, drug testing, protocol optimization, and therapeutic approaches.

PAX7 is an important regulator of stem cell maintenance conserved in several species and is widely used as a SC marker (Seale et al., 2000; Zammit et al., 2006). By PAX7 IHC, we examined the number of residing SC in muscle sections of DMD and WT piglets and detected significantly higher amounts of subsarcolemmal PAX7+ cells in DMD tissue compared to controls. This is consistent with other studies where higher levels of SC were observed in muscle tissue of *mdx* mice (Reimann et al., 2000) as well as in human DMD patients (Kottlors & Kirschner, 2010). Dystrophin is implied to be involved in cell polarity formation by binding MARK2, an essential component of cell polarity. Dystrophin-deficient SC fail to establish a proper apico-basal orientation of regulatory proteins, which affects asymmetric cell division. In turn, this favors stem cell self-renewal over the production of progenitor cells for muscle restoration thus explaining the increased amounts of PAX7+ SC along with a reduced regenerative capacity observed in DMD muscle (Dumont et al., 2015; Filippelli & Chang, 2021). Restoring cell polarity as a potential therapeutic intervention could promote asymmetric division in SC and enhance muscle repair in DMD patients (Dumont & Rudnicki, 2016; Wang et al., 2019). Interestingly, in our study, we found a similar increase in PAX7+ SC in *INT* and *PECT* muscle sections of DMD piglets compared to controls. The intercostal muscle is crucial for respirational functions whereas the pectoral muscle is mainly responsible for movement of the upper limbs (Baig & Bordoni, 2024; Lo Mauro & Aliverti, 2016). In this regard, it seems that the mechanisms in life-sustaining muscles are similar to those for locomotion, and treatment strategies to support muscle regeneration could act on several muscle groups.

In muscle tissue, dystrophin mechanically connects the cytoskeleton’s F-actin filaments to ECM proteins thus contributing to stability and integrity of the cell membrane (Ervasti, 2007). In that way, it has been hypothesized that dystrophin alters focal adhesion tension and successively is involved in force transmission and mechano-signaling between the cell and its surroundings (Gao & McNally, 2015). By applying TFM, we found that dystrophin-deficient SC generated significantly higher strain energy levels and average traction forces. In a recent study, a similar observation was made on immortalized cells from human DMD patients which display higher traction forces than their healthy counterparts (Hofemeier et al., 2022), highlighting the relevance of our porcine model for translational research approaches. Higher traction forces in SC could reflect a modified interaction between the cells and the ECM leading to alterations in the generation and response to mechanical signals that may influence signaling pathways related to proliferation and differentiation (Pang et al., 2023). Modified cell-ECM interaction could also influence effective adhering, which alters the mechanical properties of the tissue changing stiffness and elasticity of the muscle (Loreti & Sacco, 2022). Furthermore, tissue remodeling is considered another consequence of a modified cell-ECM interplay (Daley et al., 2008; Lu et al., 2011). Increased secretion of pro-fibrotic factors by SC (Biressi et al., 2014), which directly contributes to the eminent DMD feature of fibrosis, the extensive deposition of ECM-proteins like collagens substituting muscle with fibrotic tissue (Giovarelli et al., 2022), could therefore be linked to the altered mechanical properties observed in our study. Controversially, another recent publication reported lower focal adhesion tensions in a dystrophin-deficient C2C12 myoblast model (Ramirez et al., 2022). An explanation for these contradictory results could be the differences in stiffness of the substrate: while Ramirez et al. used 4.6 kPa gels, others and we cultivated the cells on 12 – 15 kPa gels mimicking the physiological stiffness of muscle tissue (Bruyere et al., 2019; Engler et al., 2004).

By RNA-seq analysis, we revealed a heterogeneity of transcriptional dysregulations in DMD SC. Whole expression profiles separated in a genotype- and stage-specific manner reflecting a high diversity in transcriptomes between DMD and WT SC. Amongst genes with the highest differential expression levels, we identified key myogenic transcription factors *MYF5*, *MYOD1,* and *MYOG*, which are indispensable for orchestrating myogenesis in SC (Asfour et al., 2018). In particular, in proliferating DMD SC increased levels of *MYF5,* a key factor for promoting and maintaining proliferation in muscle progenitor cells (Yamamoto et al., 2018; Zammit et al., 2004), are noteworthy. Elevated *MYF5* expression could point towards a hyperproliferative behavior of dystrophin-deficient SC that in turn perturbs the progression along the myogenic differentiation pathway. In healthy myoblasts, several sarcomeric transcripts are already expressed before fusion into myotubes to ensure proper assembly of a functional contractile apparatus in mature muscle tissue (Abdul-Hussein et al., 2012). We found upregulated genes related to sarcomere organization and contraction in proliferating DMD SC, which could indicate a compensatory mechanism to enhance muscle function. Further examinations of this finding are necessary, especially because we observed higher traction forces in DMD SC and sarcomeric components are essential for producing the contractile, force-generating units of muscle cells. On the other hand, significantly decreased transcript levels in proliferating DMD SC include hypoxia-associated factors. In healthy SC, hypoxia enhances differentiation in a HIF1α-dependent manner (Lundby et al., 2009; Nguyen et al., 2023). Alterations in transcript levels of hypoxia-related genes thus could hamper this response and impair myogenesis in dystrophin-deficient SC.

Gaining deeper insights into the molecular consequences of dystrophin-deficiency through proteome analysis, we found a clear separation of the WT DIFF cells from the rest of our cohort, indicating a highly impaired differentiation behavior of the DMD SC. The number of differentially abundant proteins in DMD DIFF almost doubled compared to the PROL counterpart further emphasizing the fundamental impact of dystrophin-deficiency on protein composition during myogenesis. In proliferating DMD SC, focal-adhesion-associated integrins ITGB1, ITGA7 and ITGA6 were more abundant than in the corresponding WT SC. Integrin-based adhesion sites are largely responsible for force transmission between intra- and extracellular compartments (Sun et al., 2016) and co-localize with the dystrophin-glycoprotein complex at the sarcolemma (Anastasi et al., 2003). It has been suggested that the physical proximity of the two systems implies a mutual regulation of transmitted forces and signals (Wilson et al., 2022). In this way, increased integrin expression could be an explanation for the elevated traction forces observed in DMD SC in this study. In a recent publication, it was demonstrated that isoforms of the ALDH enzyme family are essential for myogenic differentiation (Steingruber et al., 2023). In our analysis, we detected reduced amounts of ALDH1A1 and ALDH1A3 in differentiated DMD SC. This reduction could contribute to the disturbed differentiation behavior in dystrophin-deficient SC and should be validated in future investigations.

Using MOFA, a technique that allows the combined analysis of several omics data sets to explain the variability observed across samples (Argelaguet et al., 2018), we captured multi-layered effects of dystrophin-deficiency on SC. We detected an overall slightly higher contribution of the transcriptomic than the proteomic data to the observed variance between WT and DMD SC. The lower proteomic contribution could be explained by reduced detection sensitivity and incomplete sequence coverage as well as posttranslational modifications, which remain undetected (Ladakis et al., 2024). In total, we identified five factors of which factors 1 and 2 explain the majority of variance. Factor 1 showed a strong correlation with the genotype of the samples separating DMD from WT SC, the transcriptomic and proteomic signatures contributed almost equally. Factor 2 distinguished the samples based on their developmental state with a higher contribution of the transcriptomic data to the observed variance.

To our knowledge, this is the first co-evaluation and integration of the transcriptomic and proteomic data of dystrophin-deficient SC derived from a translational porcine model of DMD. We showed that lack of dystrophin highly affects a plethora of aspects during myogenesis already coming into play at early developmental stages. We demonstrated that dystrophin-deficiency has functional consequences leading to higher forces exerted by the affected SC at a single cellular stage when they are still unexposed to contraction forces. Together, our findings support the hypothesis that intracellular deficits emerging from dystrophin-deficiency account for SC dysfunctions in DMD. Future studies could include SC derived at different age stages to longitudinally monitor disease progression to identify crucial molecular events for precisely timed treatment interventions. Lastly, our dataset represents a valuable basis for perspective investigations aiming to elucidate the functions and potential therapeutic targets amongst the identified genes and proteins.

## Material and Methods

### Ethical Statement

All animals used in this study were maintained and sacrificed according to protocols approved by the responsible welfare authority (Government of Upper Bavaria; permission ROB.55.2-2532.Vet_02-19-195). All experiments were conducted in accordance with the German Animal Welfare Act and Directive 2010/63/EU on the protection of animals used for scientific purpose.

### Tissue Collection

Muscle tissue was procured freshly from 3-day-old male DMD and WT piglets immediately after euthanasia. Upon dissection of overlying tissues, samples of roughly 1.0 cube length were taken from *intercostalis* (*INT*) and *pectoralis* (*PECT*) muscles under sterile conditions. The pieces were dipped into 70% ethanol and washed with Dulbecco’s phosphate-buffered saline (PBS, GIBCO Thermo Fisher, Schwerte, Germany) supplemented with 1% Penicillin-Streptomycin (PenStrep, GIBCO Thermo Fisher) (PBS+) before being further chopped with a sterile scalpel blade. Fragments of 2-3 mm diameter were stored at 4°C in DMEM GlutaMax without Pyruvate (GIBCO Thermo Fisher) supplemented with 20% Fetal Bovine Serum (FBS, GIBCO Thermo Fisher) and 1% PenStrep (DMEM+) until the isolation was performed.

### Isolation of primary porcine satellite cells

#### Dissociation of porcine muscle tissue

Working under a sterile laminar flow, muscle biopsies were further minced using a scalpel blade. Attaching connective tissue was grossly removed. Next, the tissue was sterilized again using 70% ethanol, rinsed with PBS+, transferred to a tube containing PBS+ with 1% 1 M HEPES buffer (Sigma Aldrich, Darmstadt, Germany) solution, and centrifuged. The supernatant (SN) was transferred to a new tube and stored on ice (=SN1) until usage. For enzymatic digestion, a protease solution containing 1.5 mg/mL Protease *Streptomyces griseus* (Sigma Aldrich, 3.5 U/mg), PBS, 1% PenStrep, and 1% 1 M HEPES buffer was added to the tube to completely cover the tissue for a 1 h incubation period at 37°C. The mixture was agitated by hand every 10-15 min. After the digestion step, the tissue was centrifuged for 5 min at 400 g and the supernatant (=SN2) was added to SN1. To further dissociate the cells, the mixture was incubated with a collagenase solution consisting of 1 mg/mL Collagenase *Clostridium histolyticum* (Sigma Aldrich, 800 U/mg), DMEM without Pyruvate GlutaMax, 5% FBS, and 1% PenStrep for 1 h at 37°C, agitated by hand every 10-15 min. During the second digestion period, SN1 and SN2 were further processed: the suspension was centrifuged for 5 min at 400 g, the supernatant was discarded and DMEM+ was added. The solution was sifted through a 70-µm nylon strainer (Greiner, Frickenhausen, Germany) and stored on ice until further use (SN3). The digested tissue suspension was filtered through a 70-µm nylon strainer, centrifuged for 10 min at 400 g and the cell pellet was resuspended using SN3. The solution was sifted through a 40-µm nylon strainer (Greiner), PBS+ was added to the filtered suspension and the mixture was centrifuged for 5 min at 300 g. The cell pellet was resuspended in DMEM+ and again centrifuged 10 min at 400 g. The isolated cells were resuspended in DMEM+ and prepared for subsequent purification procedure.

#### Magnetically activated cell sorting (MACS)

The heterogeneous cell solution was further processed by Magnetically activated cell sorting (MACS) using an indirect labeling approach and two sequential separation steps. First, for depletion of unwanted cells, polyclonal rabbit anti-CD31 (1:200, ab28364, Abcam, Cambridge, UK) and polyclonal rabbit anti-CD45 (1:200, ab10558, Abcam) antibodies were added to the cell suspension in order to bind surface antigens of endothelial respectively hematopoietic cells and incubated for 30 min on ice. Subsequently, magnetically conjugated anti-rabbit IgG(-) secondary antibodies (1:15, Miltenyi Biotec, Bergisch Gladbach, Germany) were added and incubated for 30 min on ice. Next, a washing step was performed, and the pellet was resuspended in MACS buffer (Miltenyi Biotec). The MACS separator (Miltenyi Biotec) was placed under the sterile working bench and a LS column (Miltenyi Biotec) was set up on the magnet board with a tube underneath to collect the flow-through. The column was pre-balanced by rinsing with MACS buffer, the cell suspension was transferred into the column and the solution was gently sifted through by inserting the plunger. The undesired cells were retained in the column by magnetic forces and thus were eliminated from the cell suspension, whereas unlabelled cells flowed through. To target the skeletal muscle SC in the remaining solution primary antibodies for monoclonal mouse anti-Integrin-β1 (1:200, ab30388, Abcam), monoclonal mouse anti-NCAM1 (1:200, ab9018, Abcam) and monoclonal mouse anti-M-cadherin (1:200, sc-374093, Santa Cruz) and the magnetically conjugated secondary anti-mouse IgG(-) antibodies (1:15, Miltenyi Biotec) were added and incubated for another 30 min on ice. Again, a washing step was performed before transferring the solution into a MS column (Miltenyi Biotec) attached to the magnetic board with a tube placed underneath. The column was pre-balanced with MACS buffer and the solution was gently filtered through the column by using the plunger. During this separation step, the desired, magnetically labeled cells remained inside the column due to magnetic forces. Therefore, the flow-through was discarded, a new tube was placed underneath the column, and the tube and column were removed from the magnetic board together. Now, the target cells were eluted by flushing the column with MACS buffer twice. The cell suspension was centrifuged for 3 min at 400 g and the cell pellet was resuspended in growth medium consisting of Ham’s F10 Medium (GIBCO Thermo Fisher), 20% FBS, 1% PenStrep and 10 µg recombinant human basic Fibroblast Growth Factor (bFGF, PEPro Tech, Cranbury, New Jersey, USA). The purified cells were then counted with a Neubauer Chamber and directly seeded on a 10% Matrigel (VWR, Rednor, Pennsylvania, USA) -coated 12-well plate.

### Primary Cell Culture

After MACS sorting, the cells were incubated in growth medium for 3 days at 37°C with 5% O_2_ uninterruptedly, and afterwards the medium was exchanged every second day. By reaching 70-80% confluency, cells were splitted using Trypsin/EDTA 0.25% (GIBCO Thermo Fisher) and seeded into 10% Matrigel-coated culture dishes at a concentration of 5000 cells/cm^2^ for all following passages. After passage 3, cells were cultivated in 50 µg/mL collagen-coated culture dishes using Collagen I, rat tail (GIBCO Thermo Fisher). Differentiation into myoblasts was induced by nutrient starvation using Ham’s F10 Medium supplemented with 1% PenStrep and 5% Horse Serum (Biozol, Eching, Germany) for 7 consecutive days, during which medium was exchanged every second day.

### Traction Force Microscopy

#### Silicone substrate preparation

Employed 60 x 24 mm glass coverslips (Marienfeld, Lauda-Königshofen, Germany) underwent a thorough cleaning process in three steps. Initially, coverslips were sonicated in a 5% v/v solution of Hellmanex III (Sigma Aldrich, Merck) for 10 minutes, followed by sonication in 1 M potassium hydroxide for another 10 minutes and, lastly, in 100% ethanol. During each step, the coverslips received a rigorous washing in double distilled water for 5 minutes. Finally, compressed nitrogen was used to dry the coverslips. Polydimethylsiloxane (PDMS) substrates were fabricated by combining a 9:10 weight ratio of two components Cy-A and Cy-B from DOWSIL™ CY 52-276 Kit (Dow, Midland, Michigan, USA) resulting in a substrate stiffness of approximately 12 kPa. The gel solution was mixed thoroughly, centrifuged, and degassed before pipetting 350 µl of the solution onto a glass coverslip. The solution was spin-coated for 90 seconds using a home-made spinner, with acceleration and deceleration occurring every 30 seconds, to create a smooth, uniform surface. The gels were cured at 70°C for 30 min, after which they were stored at RT. To functionalize the silicone surface, the gels were treated with a solution of 7% (vol/vol) 3-aminopropyl trimethoxysilane (APTES) in pure ethanol for 2 hours. The gels were washed with pure ethanol, a 1:1 volume ratio of ethanol and PBS, and finally with only PBS, each for three cycles. As a cell culture and imaging chamber, a bottomless 8-well sticky-slide (Ibidi, Gräfelfing, Germany) was mounted on top of each PDMS substrate. After washing the chambers three times with PBS, 100 µL solution of 100 nm, red-orange fluorescent (540/560) FluoSpheres carboxylate-modified microspheres (Invitrogen Thermo Fisher) at a dilution of 1:1000 in PBS was mixed with a final concentration of 100 µg/mL 1-ethyl-3-(3-dimethylaminopropyl)carbodiimide (EDC) and pipetted onto the top surface of the gel containing 200 µL PBS. The solution was incubated for 20 min at RT and washed three times with PBS. Next, the gels were coated with a solution of 50 μg/mL collagen I with 100 μg/mL of EDC in PBS, and incubated for 2 hours at RT. After incubating, the chambers underwent multiple washing steps, initially with PBS and then with the cell culture growth medium, before being seeded with cells.

#### Traction force measurements and data analysis

To investigate differences in traction forces exerted by the dystrophin-deficient and healthy control satellite cells, primary porcine SC derived from 3-day-old DMD and WT piglets were seeded onto the PDMS substrates containing 100 nm fluorescent microspheres at a density of 5000 cells/cm^2^ in growth medium and cultivated under standard conditions for 48 hours before performing TFM. Fluorescent images of the beads embedded in the PDMS gel were captured at 40x with an inverted IX83 microscope (Olympus, Japan) equipped with a camera (XM10, Olympus) before and after trypsinization the cells using 0.5% Trypsin/EDTA (GIBCO) to obtain pairs of traction-loaded (deformed gel with cells exerting forces onto the substrate) and traction-free (relaxed gel without cells) bead images. We performed further processing and analysis in MATLAB with our customized script. The entire analysis workflow includes bead detection and tracking, traction field reconstruction, maximum traction, and strain energy statistical analysis. Fluorescent beads were detected by *detectMinEigenFeature*, a built-in point detector in MATLAB. Depending on the quality of the image, a difference-of-Gaussian filter with a standard deviation of 1.5 pixels and 2 pixels was applied to suppress background noise and enhance the bead profiles before detection. Bead displacements were then measured by optical flow tracking which is implemented by *vision.pointTracker* as suggested by (Holenstein et al., 2017). The tracking window was set to 61 pixels to acquire smooth results. Once the displacement field was generated, traction forces were calculated using Regularized Fourier Transformation Traction Cytometry provided by (Huang et al., 2019). All experiments used a constant regularization parameter 2.47*10-5 Pix^2^/Pix^2^ and the Poison ratio was set to 0.5. For analysis, we collected the maximum traction magnitude within the cell region and strain energy stored in the gel in the current field of view for all experiments and classified the results by different treatments. The TFM workflow and analysis script can be found at https://github.com/J-jianfei/TFM_workflow.

### Immunohistochemistry

Formalin-fixed and paraffin-embedded muscle tissue sections from *INT* and *PECT* biopsies derived from male DMD and WT were used for immunohistochemistry (IHC) according to standard avidin-biotin peroxidase complex method. After dewaxing and antigen-retrieval, sections were incubated overnight (o/n) at 4°C with the following primary antibodies: monoclonal mouse anti-PAX7 (1:150, Abcam, ab199010) or polyclonal rabbit anti-Dystrophin (1:100, Abcam, ab15277). The next day, sections were incubated with a biotinylated goat-derived anti-mouse IgG antibody (1:1000, Vector Laboratories, Newark, New Jersey, USA) respectively anti-rabbit IgG antibody (1:1000, Vector, BA-1000-1.5) for 1 hour at RT. Subsequently, sections were incubated with a horseradish peroxidase-labeled avidin/biotin complex (1:100, VECTASTAIN Elite ABC-HRP Kit, Vector, PK-6100) for 30 min at RT. To visualize immunoreactivity, ImmPACT DAB Substrate Kit (Vector, SK-4105) with 3,30-diaminobenzidine tetrahydrochloride dihydrate (DAB) was applied to the sections (brown color). Hemalaun was used for nuclear counterstaining (blue color). For negative control, primary antibodies were omitted and replaced by diluent. Slides were investigated via bright field microscopy using a Zeiss Axiophot® equipped with a CCD camera supported by pylon viewer. For PAX7 quantification, stained sections were scanned with a high-throughput slide scanner (Aperio AT2, Leica Biosystems) at 40x and images were taken with Aperio ImageScope (v.12.4.0.7018, Leica Biosystems). Five randomly chosen fields of view with a dimension of approximately 825 x 450 µm were analyzed by manual counting of PAX7+ cells.

### Immunofluorescence

For immunofluorescence (IF), cells were seeded in a 50 µg/mL collagen-coated 12-well plate containing a coverslip (Neolab, Heidelberg, Germany) and cultivated until reaching 70-80% confluency. Staining was performed in accordance to standard protocols. Briefly, cells were fixed with 4% formalin for 30 min at RT. Next, cells were washed with PBS and blocked with 1% BSA (Carl Roth) and 2.5% NDS (Sigma Aldrich) for 30 min at RT to avoid non-specific binding. The samples were incubated o/n at +4°C with the following primary antibodies diluted in 1 % BSA solution: monoclonal mouse anti-Desmin (1:250, Agilent DAKO, Santa Clara, California, USA, M0760) or monoclonal mouse anti-PAX7 (1:150, Abcam, ab199010). The next day, cells were washed with PBS and to reveal primary antibody binding, samples were incubated with the fluorophore-conjugated secondary antibody Alexa Fluor 488 donkey anti-mouse IgG (Invitrogen Thermo Fisher) diluted 1:500 in 1% BSA solution for 30 min at RT. Nuclei were counterstained with Hoechst 33342 (Thermo Fisher) at 1:10°000 for 10 min at RT. Coverslips were mounted on object glass slides (Topfrost) using Aqua/Poly-Mount (Polyscience, Warrington, Pennsylvania, USA). Imaging was performed with the Axio Imager Z1.m (Zeiss) equipped with an HBO Lamp (Zeiss) and AxioVision software (Zeiss, Version SE64 Rel. 4.9).

### ImmunobloUng

Total cell protein extracts were generated with radioimmunoprecipitation assay buffer (RIPA) containing 1% of 100X Protease/Phosphatase Inhibitor (Cell Signaling) according to standard protocols. The protein concentration of each sample was determined by Bradford Assay using the Protein Assay Dye Reagent Concentrate (BioRad, Hercules, California, USA) as described by the manufacturer and absorbance was detected with the Infinite F200 Pro plate reader (Tecan, Männedorf, Switzerland) at 595 nm. 25 µg of protein were separated by 6-12% SDS-PAGE and subsequently transferred to polyvinylidene difluoride (PVDF) membranes (Merck Millipore Ltd.). Next, membranes were blocked using either 1X Roti Block (Carl Roth, Karlsruhe, Germany) or 5% non-fat dry milk (Carl Roth) in Tris-buffered saline (TBS) with 0.5% Tween-20 (Carl Roth) and incubated o/n at 4°C with the following primary antibodies: monoclonal mouse anti-NCL-DYS1 (1:100, Leica Biosystems), monoclonal mouse anti-NCL-DYS2 (1:100, Leica Biosystems) and monoclonal rabbit anti-β-Actin (1:1000, Cell Signaling, Danvers, Massachusetts, USA, 4970S). The next day, membranes were incubated with horseradish-peroxidase-conjugated secondary antibodies goat anti-mouse IgG (1:5000, Abcam, ab6789) or goat anti-rabbit IgG (1:3000, Abcam, ab6721) for 45 min at RT. For visualization of protein bands, membranes were incubated with SignalFire ECL reagent (Cell Signaling) as indicated by the manufacturer, and the signal was detected with Amersham Imager 680 (GE Healthcare Amersham, Bioscience, Freiburg, Germany). Semi-quantification of immunoblots was performed using ImageJ using β-Actin signal as loading control.

### RNA isolation from cell lines, library preparation, sequencing, and data analysis

Total RNA from SC derived from 3-day-old DMD and WT animals was extracted using the NEB Monarch RNA Miniprep Kit. The protocol was performed in accordance with the manufacturer’s instructions. To recapture proliferation (PROL) conditions, samples were generated at p6 after cultivation in normal growth medium. For the differentiation condition (DIFF), samples were collected after cultivation in differentiation medium containing reduced serum amounts for 10 days at p6. To quantify gene expression in extracted RNA samples, the protocol of 3’ mRNA-seq was adapted (Bagnoli et al., 2018; Picelli et al., 2013; Picelli et al., 2014). 25 ng of total RNA was used and each sample was analyzed as a technical duplicate. To generate libraries, polyA+ RNAs were selected during cDNA synthesis through annealing to a polydT oligo containing an additional unique molecular identifier (UMI) and a well-barcode which allowed early pooling of all samples directly after cDNA synthesis. Second strand synthesis was done by RT-PCR using a template switch oligo with an integrated Illumina adapter sequence. After purification and exonuclease°I treatment, the library pool was amplified by PCR and purified again. To achieve optimal fragment lengths, NexteraXT DNA library prep Kit (Illumina, San Diego, California, USA) was used. With the Nextera transposon DNA was tagmented, a process that fragments and tags DNA with adapter sequences. For amplification of the library, a 3’ enrichment PCR was performed during which i7 and i5 adapters were attached. After purification, the quantity of the library pool (Qubit Fluorometer 4.0, Thermo Fisher) and average fragment size (LabchipGX Touch 24, Perkin Elmer/Revvity, Waltham, Massachusetts, USA) were measured. The resulting library was sequenced on a NextSeq550Dx using a 75-cycle high output kit (Read 1:16°cycle – UMI + barcode; Read 2: 76°cycles – Insert). RNAseq reads were demultiplexed using the program bcl2fastq provided by Illumina. R version 4.1.2 was used for all further analysis. Quality control was done using the command line tools fastQC (Andrews, 2010) and multiQC (Ewels et al., 2016). Umi-tool dedup (Smith et al., 2017) was used for deduplication, and the reads were aligned to the *Sus scrofa* reference genome susScr3 using STAR (Dobin et al., 2013) aligner version 2.7.10b. The number of reads mapping to each gene was also obtained from STAR using the parameter quantMode GeneCounts. edgrR (Robinson et al., 2010) was used for differential expression analysis and gene set enrichment analysis was done using fgsea (Korotkevich et al., 2021). The RNA-seq data discussed in this publication have been deposited in NCBI’s Gene Expression Omnibus (GEO) (Edgar et al., 2002) and are accessible through GEO Series accession number GSE287756 (https://www.ncbi.nlm.nih.gov/geo/query/acc.cgi?acc=GSE287756).

### Sample preparation for LC-MS/MS and proteome data analysis

For proteome analysis, SC from 3-day-old DMD and WT animals were harvested by Trypsin/EDTA 0.25% incubation followed by 5 washing steps with DPBS + 10% PenStrep. Similar to the approach for RNAseq, cell samples were harvested at two different time points in order to investigate protein expression in PROL and DIFF states: PROL samples were collected at p3 and p6, DIFF samples at p4 and p7. Samples were stored at -80°C until LC-MS/MS measurements. Using ultrasonication (Sonopuls GM3200, BR30, Bandelin, Berlin, Germany) cells were then lysed in 8 M urea/0.4 M NH_4_HCO_3_. Protein concentration was assessed photometrically (Pierce 660 nm, Thermo Fisher Scientific). Samples were reduced with dithioerythritol at a final concentration of 5 mM at 37°C for 30 min and alkylated with iodoacetamide at a final concentration of 15 mM at ambient for 30 min. Protein was digested with lysyl endopeptidase (1:100, enzyme : protein, FUJIFILM Wako Chemicals Europe, GmbH, Neuss, Germany) at 37°C for 4 hours, diluted to 1 M urea with water, and then digested at 37°C overnight after adding trypsin (1:50, enzyme : protein, Promega, Fitchburg, Wisconsin, USA). Samples were then dried and resuspended in 0.1% formic acid in water. For mass spectrometry analysis, 1 µg of peptides were separated and analyzed using an Ultimate 3000 RSLC chromatography system coupled to a QExactive HF-X mass spectrometer (both Thermo Fisher Scientific). For separation, peptides were first trapped on a PepMap 100 C18 trap column at a flow rate of 5 µL/min of 0.1% formic acid in 1% (V/V) acetonitrile in water. Peptides were then separated with an Easy-Spray column (75 µm x 50 cm; 2 µm; both Thermo Fisher Scientific) at a flow rate of 250 nL/min with eluent A 0.1% formic acid in water and eluent B 0.1% formic acid in acetonitrile using a multi-step gradient: initial ramp from 3% eluent B to 6% in one minute followed by an 80 min ramp to 20% followed by a 9 min ramp to 40%. MS spectra were acquired with a top 15 data-dependent method. For protein identification, MaxQuant (v. 2.0.3.0) (Cox & Mann, 2008) and the porcine subset of the RefSeq database were used (retrieval date: 05.10.2022). The mass spectrometry proteomics data have been deposited to the ProteomeXchange Consortium via the PRIDE (Perez-Riverol et al., 2025) partner repository with the dataset identifier PXD059983.

### Multi-omics factor analysis

Multi-omics Factor Analysis (MOFA) was used to integrate the RNAseq and proteomics datasets. MOFA identifies latent factors (LFs) across different -omics datasets referred to as views. MOFA takes M data matrices as input (Y_1_, …, Y_M_), each representing a different data modality, with co-occurring samples that may have unrelated features and varying quantities. It decomposes these matrices into a matrix of latent factors (Z) for each sample and M weight matrices (W_1_, …, W_M_), one for each data modality (Signorelli et al., 2023).

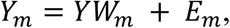

where Z is a matrix of LFs shared across all views, W_m_ is a matrix of view-specific factor loadings that quantify the contribution of each molecule in view m to each LF, and E_m_ is a matrix of error terms (Signorelli et al., 2023).

### Statistical Analysis

All data were analyzed with GraphPad PRISM (Version 5.03), Perseus (Version 1.6.7.0) (Tyanova et al., 2016), R (R Core Team, Version 2023.06.0) and MATHLAB (R2022b). Values are presented as mean and ± standard error mean (SEM) or standard deviation (SD). To calculate p-values, t-tests, ANOVAs, Tukey Honest Significant Differences, and two-tailed Mann-Whitney U tests were used. The threshold for significant differences was considered as p < 0.05 and significance levels are indicated as p < 0.001 (***), p < 0.01 (**), p < 0.05 (*), and p > 0.05 (n.s.). The number of analyzed animals (n) and p-values are listed in the figure legends.

## Acknowledgements

The authors would like to thank all members of the laboratory who supported the project. We thank Lisa Pichl for her support during cell isolation and staining procedures. We thank Patrizia Torelli for her help in cell isolation and cell culture work. We thank Sandra Baur for supporting general lab work related to cell culture, western blot and cell staining protocols. We thank Kathrin Simon for her assistance during traction force measurements. We thank Eva Mayr for her expert RNA sequencing service. We also thank Miwoka Köster for technical assistance during proteomics measurements. A.M. and B.S acknowledge funding from the European Research Council (ERC) under the European Union Horizon 2020 research and innovation program (G.A. 852585).

## Author contribution

S.F. conceived the study and designed the experiments with intellectual input from N.P., B.S., T.F., E.W., J.S. and K.M. M.S. and C.K. provided the samples. S.C., J.B.S, A.M. and J.J. conducted bioinformatics analysis and gave input towards data interpretation. N.P., B.S., T.F., E.W. gave conceptual advice and supported data evaluation. J.S. and K.M. supervised the project. S.F wrote the original draft, reviewed, and edited the manuscript with input and comments from all authors. All authors approved the final version.

